# 3D-Printed LEGO^®^-inspired Titanium Scaffolds for Patient-Specific Regenerative Medicine

**DOI:** 10.1101/2023.03.30.534953

**Authors:** Seunghun S. Lee, Xiaoyu Du, Thijs Smit, Elisa G. Bissacco, Daniel Seiler, Michael de Wild, Stephen J. Ferguson

## Abstract

Despite the recent advances in 3D-printing, it is difficult to fabricate implants that optimally fit a defect size/shape. There are some approaches to resolve this issue, such as patient-specific implant based on CT images, however, it is labor-intensive and costly. Especially in developing countries, affordable treatments are required, while still not excluding these patient groups from manufacturing advances. Here, a SLM 3D-printing strategy was used to fabricate a hierarchical, Assemblable Titanium Scaffold(ATS), which can be manually assembled in any shape or size with ease. A surgeon can create a scaffold that would fit to the defect right before the implantation during the surgery. Additionally, the direct inclusion of micro- and macroporous structures via 3D-printing, as well as a double acid-etched surface treatment(ST) in the ATS, ensure improved nutrient flow and cellular activity. Different structures were designed, 3D-printed and then surface treated for the ST groups. Both individual and stacked ATS have sufficient mechanical properties to withstand physiological loading, and the porous groups resulted enhanced cell proliferation, mineralizaton and osteogenesis compared to non-porous group. Furthermore, successful cell attachment and migration between the assembled ATS were observed. Finally, we demonstrate possible medical applications that reveal the potential of the ATS through assembly.

## 1. Introduction

Over the past decades, as the number of bone fractures and orthopedic-related pathologies has increased with the exponential growth of the elderly population, bone tissue engineering (BTE) has emerged as a strategy with potential to address these problems [1–4]. Especially, additive manufacturing with various 3D-printing technologies has facilitated a significant development in scaffold design for BTE [5, 6]. However, despite these advances, it is often difficult to fabricate implants that optimally fit the defect size and complex 3D shape. There are strategies to resolve this issue, such as patient-specific scaffolds designed based on CT images of the patient, however, this process is labor-intensive and costly [7, 8]. Especially in developing countries, where access to medical imaging equipment is limited and affordable treatment options are required, it is not an ideal solution that is universally accessible [9]. Thus, the development of scaffolds that are affordable, easy to use, mechanically strong, biologically effective and can be adjusted and fit to any size and shape of defect is of key importance for a broader adoption of bone tissue engineering [10, 11].

Here, we report on an Assemblable Titanium Scaffold (ATS) system, a new 3D-printed scaffold, produced using a selective laser melting (SLM) 3D-printing technique, that can be manually assembled in any shape or size with ease. It was inspired by the features of LEGO^®^ building blocks, which allow users to effortlessly scale up/down and assemble to any complex structures with unlimited combinations. Surgeons can swiftly and intuitively create a scaffold that would fit to the defect, immediately prior to implantation during the surgery, without any need for special instruments. Titanium is the material of choice for the ATS, since the scaffold has to be mechanically strong to withstand physiological loading. Titanium is one of the most popular materials for bone implants, due to its excellent mechanical properties, proven biocompatibility with living tissues and high durability [12, 13]. Nevertheless, The osteoconductivity of titanium mainly depends on the surface chemistry and surface topography, and since the Young’s modulus of titanium is much higher than that of cortical bone, bulk titanium scaffolds can cause stress shielding [14]. To overcome these drawbacks, a double acid-etching surface treatment and interconnected porous structure design were incorporated in the ATS. Double acid-etching creates rough surfaces on titanium, which improves cellular activity like attachment, proliferation and differentiation [13, 15, 16]. Compared to the conventional acid-etching technique, which creates rough surfaces at the microscale, double etching creates nano-rough surfaces which promotes higher protein adsorption and enhances bioactivity [17, 18]. In addition, the interconnected porous structure design ensures sufficient nutrient and oxygen flow, vascularization, cell proliferation, migration and enhanced osteogenesis for the ATS [19]. Moreover, an open-porous scaffold may promote a more stable fixation and better osseointegration with the host bone due to interlocking between the scaffold surface and the surrounding tissue [20, 21]. In this study, we fabricated and compared several ATS groups, with/without surface treatment and incorporating various porous structures. We investigated their mechanical characteristics and cellular activity, including osteogenic capacity.

## 2. Results

### 2.1 3D printing of Assemblable Titanium Scaffold (ATS)

To allow pre-operative assembly and fitting to the defect, the main exoskeleton design of the ATS is similar to conventional interlocking LEGO^®^ toy blocks. Three designs were created, with different porous structures which may affect the bioactivity (Figure 1a). Three different ATS groups (non-porous: NP, semi-porous: SP, ultra-porous: UP) were 3D-printed by selective laser melting (SLM) technique, with outer nominal dimensions of 8 mm × 8 mm × 5.8 mm per unit, 200–500 μm thick struts and different pore size configurations (SP: channel size of 400 × 400 μm^2^ & 600 × 600 μm^2^ with porosity of 50% and UP: 600 × 600 μm^2^ & 800 × 800 μm^2^ with porosity of 62%) as shown in Figures 1b and S1. After 3D printing, double acid-etching treatment was carried out on native SLM ATS to fabricate surface textured (ST) ATS by creating homogenously rough nanostructures on the titanium surfaces. As shown in figure 1c, all SLM groups could be 3D-printed, without any defects. At the macro scale, SLM groups showed a silverly gray color and rough surfaces due to titanium particle deposition during SLM, while ST groups demonstrated a dark gray color and smoother surface than the SLM groups due to titanium particle removal during double acid-etching. There was no damage or loss of assembly functionality caused by double acid-etching treatment. After fabrication, the square protrusion on the top of the ATS interlocked precisely with the concave region in the bottom of the mating part by simply positioning one above the other (Figure 1d). We were able to assemble all ATS groups in any shape or size with ease. As shown in figure 1e, the scaffolds could be assembled in a multistacked form with various dimensions, for example forming a stacked implant for a random size defect, a pyramidal shape and an implant similar to a conventional spinal fusion cage. Numerous click-assembled configurations are possible with just a single base ATS design, showing the potential for low cost patient specific treatments.

**Figure 1.**
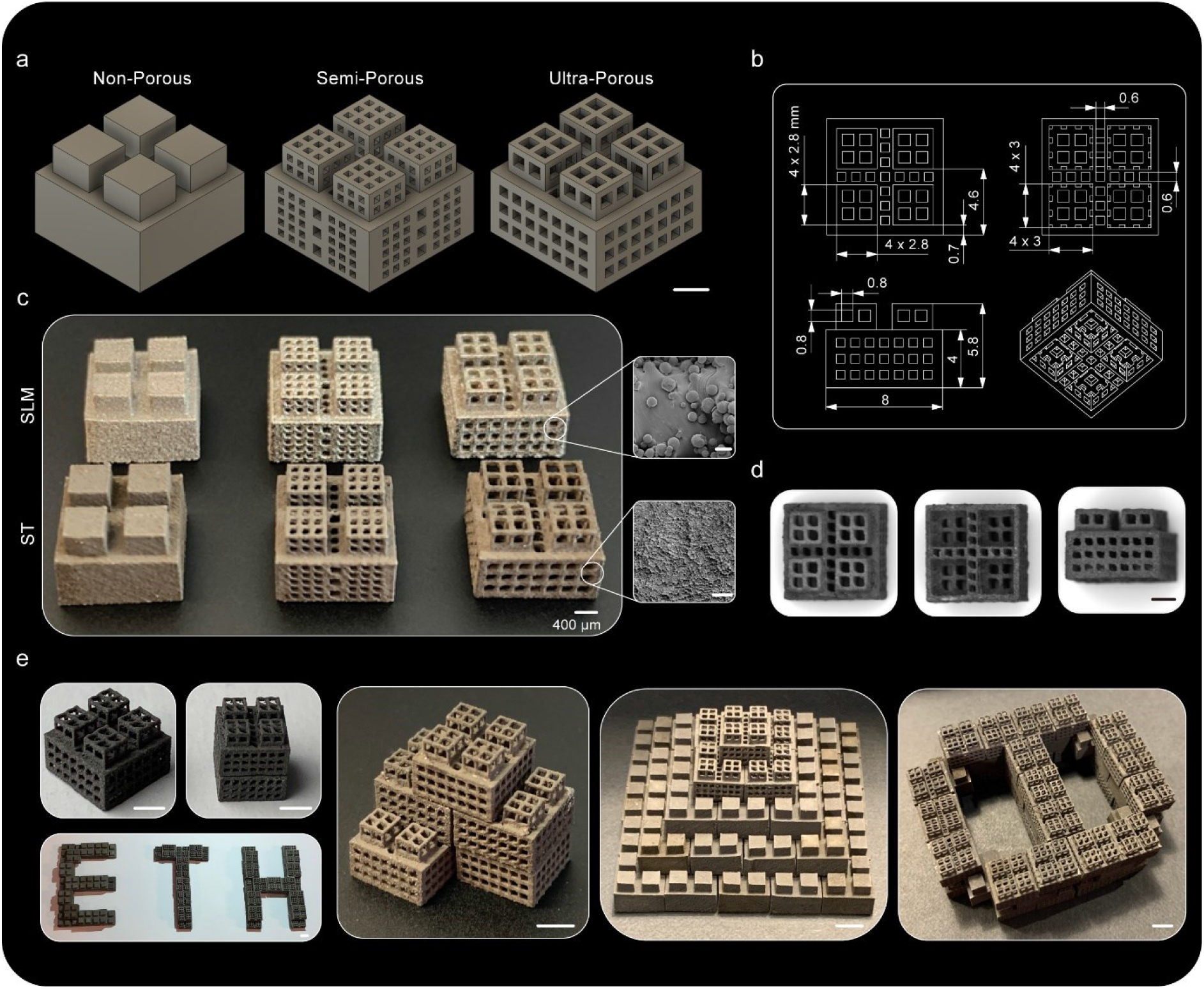
Design and Fabrication of Assemblable Titanium Scaffold (ATS) **(a)** Orthogonal projection of 3D designs of single ATS unit with different structures: Non-Porous (NP, left), Semi-Porous (SP, middle) and Ultra-Porous (UP, right). Scale bar: 2 mm. **(b)** Representative design of single UP ATS scaffold with dimensions and different perspectives. **(c)** Representative photographs of selective laser melting (SLM, top) and double acid-etched (ST, bottom) ATS with different structures NP, SP and UP. Representative SEM image in the dashed line demonstrates the surface of SLM and ST treated ATS scaffolds (Scale bar: 50μm and 25μm for top and bottom SEM respectively). **(d)** Representative microscopic images of ST treated UP ATS scaffold with different perspectives (top view, left), (bottom view, middle) and (side view, right). (a-d) Scale bar: 2 mm. **(e)** Representative photographs of ATS scaffolds assembled in various forms. Scale bar: 4 mm.

### 2.2 Surface characterization, Mechanical Durability and FEA of ATS

After fabrication, the inner structure and surface topology of the ATS was investigated, as essential factors governing cell response and osteogenesis. As shown in the figure 2a and S2, the longitudinal and the transverse cross-section of SP and UP ATS demonstrate well-interconnected porous structures which could promote easier cell migration, vascularization and nutrition inflow for both a single ATS unit or assembled ATS systems. A micro-CT scan of the ST UP ATS scaffold allowed us to observe the interconnected pores in both lower and higher transverse cross-section region of the scaffold (Fig. 2b). The surface topology difference between SLM ATS and ST ATS scaffolds was compared using field emission scanning electron microscopy (SEM). As shown in Figures 2c and S3, a smooth surface with heterogeneously spread titanium microparticles was observed on SLM groups while the surfaces were homogenously nanotextured on ST groups due to the double acid-etching treatment. As expected, the 3D topographic images of the surface (283 × 283 μm^2^) captured by confocal laser scanning microscopy (CLSM) demonstrated that the SLM group had higher area roughness parameter, Sq, than those of ST which is also confirmed by the graphs of the quantitative roughness profiles of the regions. This is due to the titanium micro particles on SLM groups which increased the surface roughness at the macro scale. In contrast, as shown in the image and graph of roughness profiles on the ST groups, despite its lower Sq value, the ST groups showed homogenously nano-textured rough surfaces and more rough surfaces at the micro level compared to the SLM group, which confirms the successful ST treatment on titanium scaffolds.

**Figure 2.**
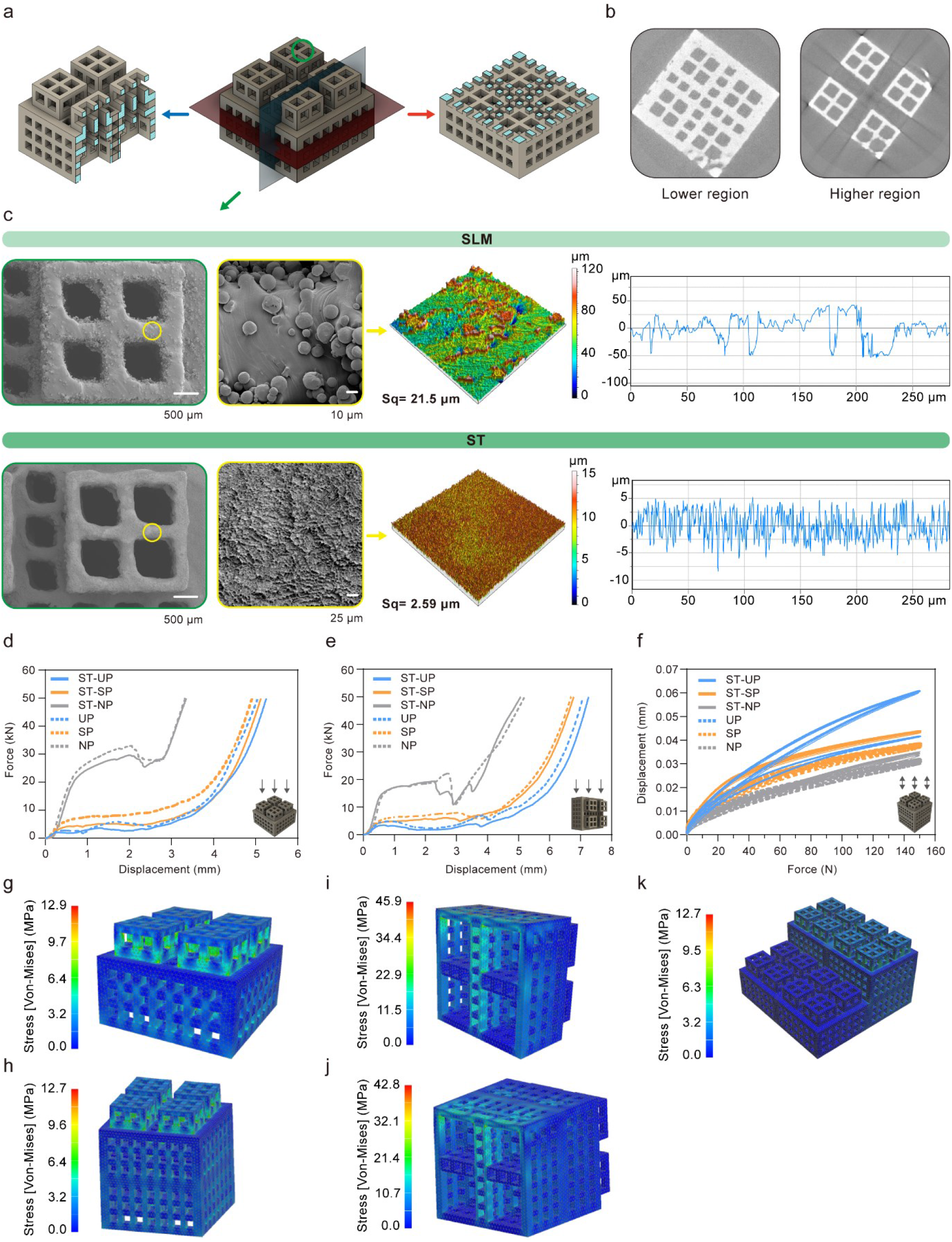
Characterization of ATS system. **(a-c) (a)** 3D diagrams showing the longitudinal (blue plane, left) and the transverse (red plane, right) cross section of UP ATS scaffold demonstrating the interconnected porous structure. **(b)** Representative micro-CT scan image of ST-UP ATS scaffold. Left image represents lower transverse cross section and right image represents higher transverse cross section of the 3D-printed scaffold. **(c)** Representative field emission scanning electron microscopy (SEM) images of SLM and ST treated UP ATS scaffolds. SEM images in yellow dashed box demonstrate the magnification of the yellow circled region on the left SEM images to show the surface structure in microscale. The representative confocal laser scanning microscopy (CLSM) 3D topographic images show the degree of surface roughness of SLM and ST scaffold surfaces, and graphs on the right represent the quantitative roughness profiles of CLSM 3D topographic images. The measured size of each sample was 283 × 283 μm. **(d-f)** Mechanical properties of ATS scaffolds. Representative force-displacement curve of ATS scaffolds under (d) vertical compression and (e) lateral compression. (f) Representative displacement-force curve of longitudinally assembled ATS scaffolds (assembled from two ATS units) under cyclic loading compression. Right bottom images of ATS with arrows represent load case. Arrows represent the direction of the loading. **(g-k)** Finite element analysis (FEA) of single and assembled ATS scaffolds under the maximum intradiscal pressure of 1.25 MPa. The pressure load was applied on the top surface of the scaffold with the bottom surface fully constraint in (g) single ATS, (h) symmetrically assembled two ATS and (k) asymmetrically assembled six ATS conditions and on the side surface with the opposite side fully constraint in (i) single ATS and (j) symmetrically assembled two ATS.

In order to investigate ATS’ mechanical properties as an implant, compression tests were performed on single ATS units. Based on the force versus displacement curves of ATS samples, both NP and ST-NP exhibit higher stiffness and strength than the other groups in both vertical and lateral compression (Figures 2d and 2e). As the porosity of ATS increased, the stiffness and strength decreased (Figure S4). Overall, ST ATS scaffolds which were treated by double acid-etching and where some material has been removed, demonstrated inferior mechanical properties compared to native SLM ATS scaffolds which was about 3.8%, 26.3% and 12.4% reduction for NP, SP and UP respectively. Note that all ATS scaffolds were ductile and no brittle fracture was observed in the tests. To confirm the mechanical stability of assembled ATS configuration, two ATS units were assembled and tested under cyclic loading to 150N (5 cyles at 0.05 Hz), a nominal compressive pressure of 2.35 MPa. As shown in figures 2f and S5, all samples maintained their integrity under dynamic cyclic loading. The NP and ST-NP groups showed the least variation in displacement, while the UP and ST-UP groups showed the greatest variation in displacement. As expected, the NP and ST-NP groups showed higher stiffness values than the other groups (Figure S4). Additionally, a hysteresis loop was observed in all samples, and the UP and ST-UP groups showed a larger hysteresis loop, indicating a higher energy dissipation.

Finite element analysis (FEA) was performed to examine the mechanical stability of the ATS in single unit and assembled unit configurations under simulated physiological loading. A distributed pressure load of 1.25 MPa, the maximum intradiscal pressure [22], was applied in the longitudinal or transverse direction. As shown in figure 2g, the maximum Von Mises stresses predicted for single loaded scaffold were well below the yield strength of the material for both the top and side load-case. The top loaded scaffold shows stress concentrations around the pores, with a maximum of 12.86 MPa. The side loaded scaffold shows higher stress concentrations compared to the top loaded scaffold. For the side loaded scaffold, the stress concentrations, with a maximum of 45.86 MPa, are in the corners where the ribs connect to the wall (Figure 2h). Assembled ATS were examined to check the difference in mechanical stability in case of assembly. As shown in Figures 2i and 2j, similar stress distributions were observed, compared to the single loaded ATS. The maximum Von Mises stresses at the locations of the concentrations are 12.74 MPa and 42.82 MPa for the top and side loaded assembled ATS, respectively. Stress concentrations of an asymmetric assembled ATS system (six ATS stacked in structure of staircase) were investigated as well to confirm its stability. The result showed a maximum stress concentration of 12.69 MPa which is similar to the single loaded case and the stress concentrations are located around the pores and in the corner of the interface between the scaffolds in the assembly (Figure 2k). Based on mechanical testing and FEA, both single and assembled system in various forms have adequate mechanical properties to withstand loading in the physiological system.

### 2.3 Cellular Activity and Migration on ATS system

*In vitro* tests were conducted to investigate the biological response and bioactivity of all SLM and ST conditioned NP, SP and UP ATS groups. After fabrication, the cell viability of pre-osteoblasts was evaluated on all testing scaffolds by live and dead assay 1 day after seeding cells. Based on the confocal laser scanning microscopy (CLSM) images, most of the cells on all the scaffolds were alive (Figure 3a). Cell viability was higher for ST groups than for native SLM groups, but overall, both SLM and ST groups demonstrated greater than 90% cell viability, suggesting all scaffolds to be biocompatible (Figure 3b).

**Figure 3.**
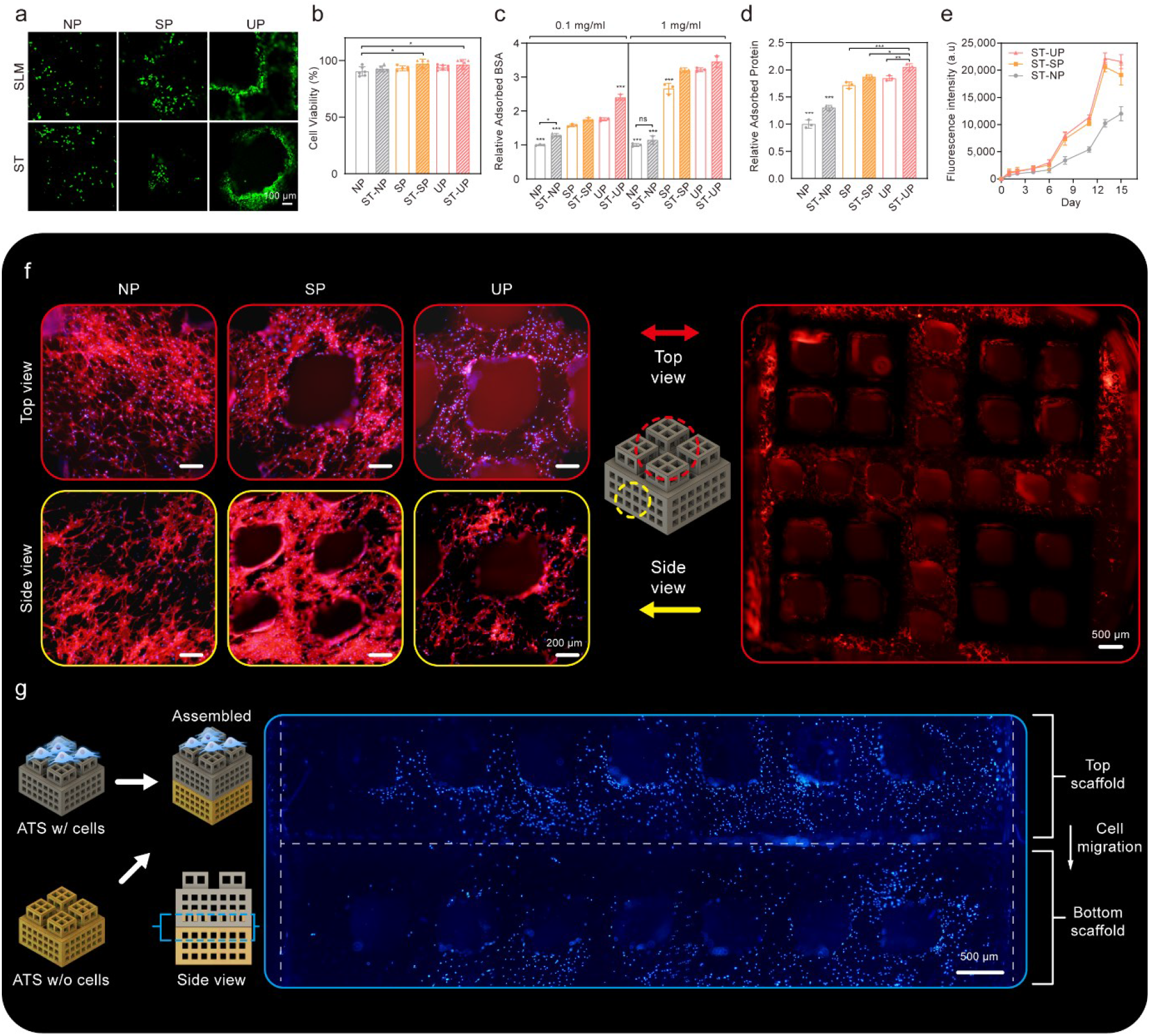
Cellular activity on ATS system. **(a)** Representative CLSM images show the live/dead fluorescent assay of pre-osteoblasts that have been seeded on SLM or ST treated NP, SP and UP ATS scaffolds. (green: live, red: dead) 1 x 10^5^ cells were seeded on scaffolds and stained by live/dead assay after 24 h of seeding. **(b)** Quantitative analysis of cell viability from live/dead assay (n=5). **(c-d)** Bioactivity analysis by protein adsorption measurement. **(c)** Adsorbed bovine serum albumin (BSA) protein amounts on the testing scaffolds, depending on 0.1 mg/ml and 1 mg/ml of BSA solution. **(d)** Relative adsorbed protein on the scaffolds when submerged in growth medium with 10% fetal bovine serum(n=3). **(e)** Quantitative analysis of cell proliferation rate in ST treated NP, SP and UP ATS groups (n=4). Error bars indicate SD. **(f)** Representative CLSM images that show the staining of actin microfilament cytoskeletal protein (red) and nuclei counterstained with DAPI (blue) of the cells after 7 days of culturing on ST treated NP, SP and UP ATS groups. Left six images show the cell attachment and morphology from the top and side view from ST-NP, SP and UP groups, and the right single image show the overall cellular attachment from the whole ST-UP. **(g)** Cell migration from top scaffold to bottom scaffold after assembly and culturing for 7 days. First, the cells were seeded onto single ST-UP ATS and cultured for 3 days. Then, the scaffold was assembled with ATS which does not have cells attached and stained with DAPI after 7 days of culturing in assembled form. In the image, blue dot represents cell nuclei, and the white dashed lines represent the boundary of the top and bottom ATS scaffolds.

The protein adsorption capacity of scaffolds was measured by submerging them in bovine serum albumin (BSA) solution to evaluate the bioactivity of the scaffolds and to determine which surface treatment groups provide a favorable cell environment through protein binding. As shown in figure 3c, non-porous NP groups presented the lowest protein adsorption among the groups while the UP, the group with higher porosity, showed a higher amount of BSA adsorbed compared to the SP, the group with lower porosity. Additionally, all ST groups demonstrated a higher protein adsorption capacity than native SLM groups due to the increase of surface area from nano-rough surfaces caused by double acid-etching. An additional protein adsorption test was also performed with growth medium containing 10% fetal bovine serum (FBS) to mimic the *in vivo* environment, and this demonstrated similar results, that the surface area increase from ultraporous structure and ST treatment led to the highest protein adsorption capacity on the ST-UP ATS scaffold (Figure 3d). As the ST groups presented overall better biocompatibility and bioactivity, the SLM groups were excluded from further *in vitro* experiments.

The proliferation rate and morphology of cells on ST-ATS groups were investigated. First, the cell proliferation rate on the scaffolds was evaluated for 15 days by presto blue assay. As shown in the figure 3e, from day 1 to day 6, there was not much difference in cell proliferation rate among the groups, but from day 8, ST-SP and ST-UP groups showed gradually higher cell proliferation rates than the ST-NP group. On day 15, ST-UP demonstrated the highest cell proliferation among the group, however, there was no significant difference to the ST-SP group. Next, for the cell morphology, cells seeded on all single ST ATS units were observed via actin and dapi staining after 7 days of culturing. As shown in both top and side views of the scaffolds, cells on all testing groups were homogenously well attached and exhibited round, polygonal morphology with distinct and thick stress fibers (Figure 3f). Interestingly, more cells were observed on ST-NP compared to ST-SP and ST-UP. Along with the proliferation result, this indicates that simultaneously, cell infiltration and migration occurred in the porous scaffolds and there was cell proliferation inside the interconnected porous structure. Additionally, the CLSM image of whole ST-UP and SEM image of cells on ST-UP demonstrate that the cells are homogenously spread and well attached not only on the outer surfaces, but also around the pores of the scaffold (Figures 3f right and S6).

After checking bioactivity and cellular activity on single ATS, cell migration between the ATS units in the case of assemblies was evaluted. First, cells were seeded on single ST-UP ATS and cultured for 3 days. Then, cell-laden ATS (top) were assembled with acellular ATS (bottom), to make stacked ATS configuration, and cell migration was observed after 7 days of culturing. As shown in figure 3g, although the cells were not homogenously spread on the bottom ATS compared to the top ATS, many cells were seen on the bottom ATS, indicating successful cell migration from the top to bottom scaffold via assembly. Interestingly, a large number of cells on the bottom of the scaffold was also observed near the pores which emphasizes that cell migration occurred via the interconnected pores in the assembled ATS units.

### 2.4 Osteogenic Effect and Potential Medical Applications of ATS

Following cellular activity, the mineralization and osteogenic effect of ATS scaffolds were evaluated. First, ARS staining was used to check the calcium deposition of pre-osteoblasts seeded on ST-ATS after culturing cells with osteogenic medium (OM) for 1 and 2 weeks. As shown in figure 4a, ST-UP demonstrated a significantly higher calcium deposition than other groups while the ST-NP showed the least calcium deposition on week 1. Similarly, on week 2, the ARS staining presented similar results as the data on week 1, with a greater difference of mineralization between the groups (Figure 4b). ST-SP and ST-UP exhibited 2.9 ± 0.10 and 3.6 ± 0.13-fold higher than ST-NP, respectively. This indicates that porous structures of ATS led to higher calcium deposition by increasing the available surface area for proteins and cells, which enhances bioactivity.

**Figure 4.**
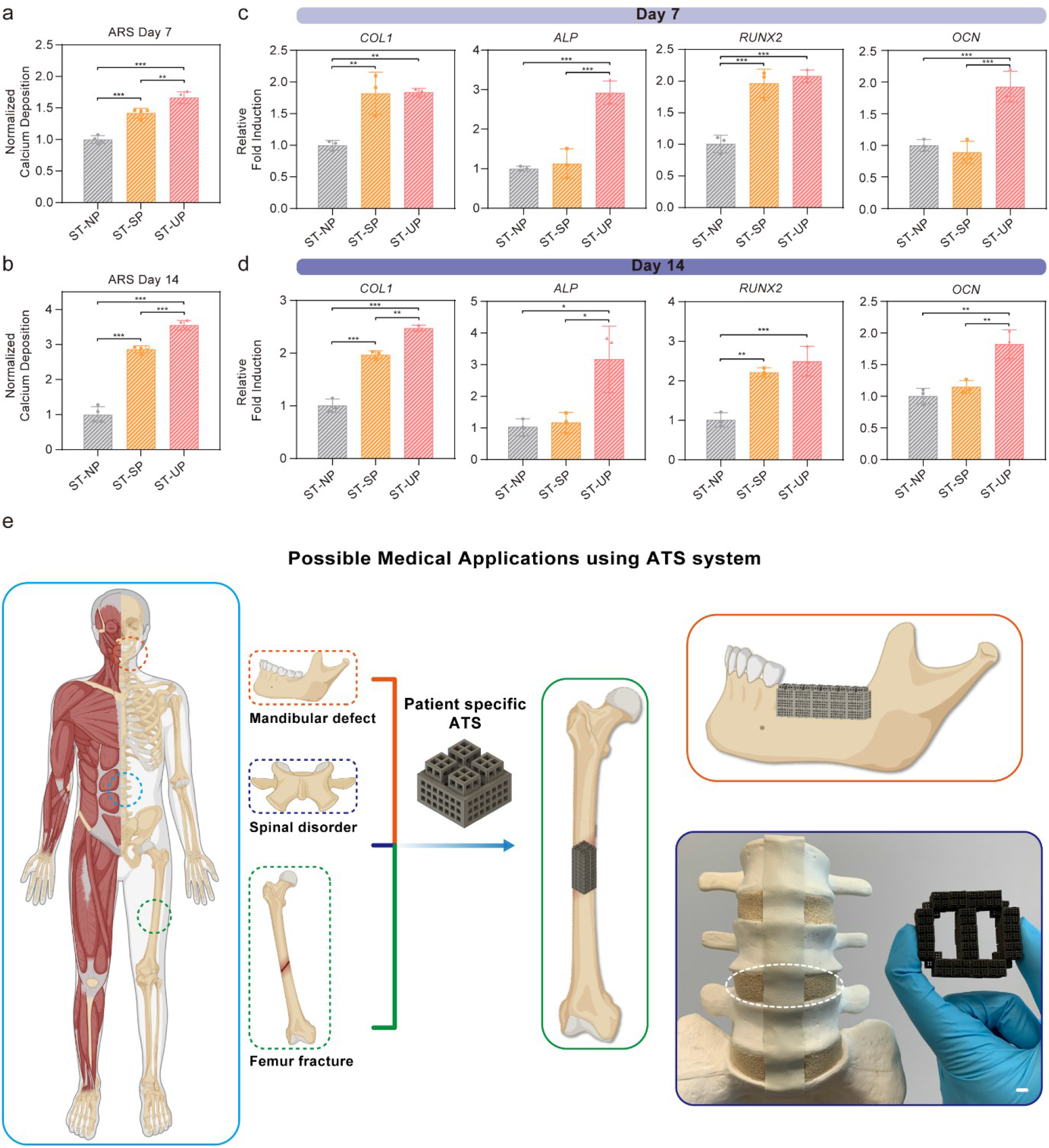
Osteogenic capacity and Potential Medical Approach as ATS system. **(a-b)** Quantitative analysis of Alizarin Red S (ARS) staining after **(a)** 7 days and **(b)** 14 days of osteogenic differentiation of pre-osteoblasts seeded on the ST-NP, SP and UP ATS groups. **(c-d)** Relative fold induction of osteogenic genes on **(c)** day 7 and **(d)** day 14. Pre-osteoblasts were seeded on the scaffolds and cultured with osteogenic medium. Error bars indicate SD (n = 3). **(e)** Schematic illustration of possible medical applications using ATS system. Three different assembled ATS implants are demonstrated as example solutions for defects in physiological system such as mandibular defect (red), spinal disorder (blue) and femur fracture (green). Scale bar: 4 mm.

ARS results were further verified by quantitative real-time PCR analysis of osteogenic gene markers. In line with ARS analysis on week 1, ST-UP exhibited the highest expression of osteogenesis marker *COL1*, *ALP*, *RUNX2* and *OCN*, with 1.8 ± 0.06, 2.9 ± 0.30, 2.1 ± 0.10 and 1.93 ± 0.25-fold higher compared to those of ST-NP, respectively (Figure 4c). Interestingly, ST-SP showed the second-highest osteogenic gene expressions, but it showed significantly greater levels than ST-NP for only two genes, *COL1* and *RUNX2*. Finally, on week 2, the result was similar as that of week 1. As shown in figure 4d, ST-UP was the group with the highest osteogenic gene expression *COL1*, *ALP*, *RUNX2* and *OCN*, with 2.5 ± 0.06, 3.17 ± 1.06, 2.5 ± 0.38 and 1.82 ± 0.23-fold higher compared to those of ST-NP, and the ST-SP showed the second highest among the groups.

As an additional proof of concept, possible medical applications were demonstrated by assembling ATS in several configurations. Possible combinations of ATS assembly are unlimited, such that they can be applied to any osteogenic defect that requires osseointegration with mechanical stability, such as vertical ridge augmentation of mandibular bones, implantation for femur fractures and spinal fusion cage for vertebrae (Figure 4e).

## 3. Discussion

In the present study, a 3D-printed LEGO^®^-inspired Assemblable Titanium Scaffold (ATS) system was developed, which can be manually assembled in any shape or size with ease and can enhance cellular activity, providing a potential low-cost patient-specific implant supporting bone tissue regeneration. Titanium scaffolds were 3D-printed using SLM, which were assigned to two groups: SLM and ST groups based on the surface treatment with double acid-etching. Also, three different structures (NP, SP and UP) were designed based on the micro and macro porosity to find an ideal scaffold that can provide a beneficial cellular environment for bone growth. From the SEM and microCT images, ATS systems were consistently well-printed, with consistent forms, and incorporating the desired porous structures that are interconnected without any major defects (Figures 2b, 2c and S3). These porous structures of the SP and UP groups are proposed to be essential, not only for cell infiltration, but also for efficient nutrient flow and vascular ingrowth for tissue healing [23, 24]. Additionally, double acid-etching on the ST groups was successful and all the surfaces of the ST groups demonstrated homogenously nanorough surfaces, unlike the SLM groups, which had residual titanium particles adhering on the surfaces. The nanorough surface of the scaffold plays an important role in improving cellular activity such as attachment, proliferation and differentiation [15, 16, 25]. Especially, the double acid-etching surface treatment creates homogenously nano-rough surfaces which allow more protein adsorption and enhances cell adhesion and proliferation [17, 18].

The mechanical stability of ATS systems was evaluated via compressive mechanical testing and FEA, confirming that all the ATS groups showed appropriate mechanical properties for use as a bone implant, due to the intrinsic stiffness and strength of the base titanium material (Figures 2d and 2e). Compared to ceramics, titanium is more ductile and avoids possible brittle fracture. In addition, the stiffness of ATS can be tailored by adjusting the porosity of ATS to match the stiffness of natural bone. This can prevent any mechanically-driven complications such as stress shielding and implant related osteopenia [14, 26, 27]. Moreover, the cyclic loading tests demonstrated the mechanical integrity of the assembled scaffold under physiological compressive loading conditions, which ensures the stability and safety of the implant even in case of complex assemblies (Figures 2f and S5). Further, stress concentrations around the pores and in corners were shown to be modest, relative to the material’s yield strength, and not significantly altered in assembled configuration, compared to the single ATS (Figures 2i, 2j and 2k).

From the *in vitro* experiments, more protein adsorption and higher cell viability were observed for the groups with higher porosity, and for the ST group compared to SLM group which was likely facilitated by the nanorough surface from ST treatment and porous structures (Figures 3a-3d). Porosity and roughness both increase the available surface area of ATS, and consequently, these provides a higher bioactivity by inducing more protein adsorption and possible apatite deposition in physiological system [28–31] [31–35]. Especially, in case of titanium based scaffold, there has been many studies that emphasized the importance of roughness and porosity for bone tissue engineering [12, 21, 36–38]. In the study of de Wild et al., surface modification of titanium using sand blasted treatment and sand blasted acid-etched treatment demonstrated significantly higher bone formation compared to native SLM in *in vivo* rabbit calbarial defect model [36]. Furthermore, Takizawa et al. confirmed the effect of rough and porous titanium scaffold by using titanium fiber plates [12]. This titanium fiber plate showed higher contact with regenerated tissue compared to convential titanium plate due to its porous and rough structure while still functioned as a conventional titanium scaffold. Therefore, although all the cells were well attached and spread on all testing groups, based on CLSM images, ST-SP and ST-UP groups exhibited a higher cell proliferation than ST-NP due to a synergistic effect from the porous structure and nanorough surface together (Figures 3e and 3f). In addition, less cells were observed on the outer surface of the porous scaffolds, ST-SP and ST-UP, than ST-NP, despite higher cell proliferation rate. This indicates that in the ST-SP and ST-UP groups, the cells attached not only to the outer surface, but also the inner surfaces and inner pores of the scaffold. These cells on the inner and outer surfaces of the scaffold will form an interconnected network form bone tissue spanning the implant. In addition, porous structure of SP and NP groups ensures high permeability as a scaffold which improves vascularization and the amount of bone ingrowth and the cell growth and the proliferation because it can provide sufficient nutrient and oxygen inflow [39, 40]. It is a same motif which was confirmed in the other studies. For instance, Kelly et al., investigated osseointegration into porous 3D-printed titanium implants covering a wide range of porosity (0%-90%) in ovine bicortical defect model, and demonstrated that bone ingrowth increases with increasing porosity which explains the potential of ATS [37]. Moreover, the larger pores with high permeability will be occluded later than the smaller pores during tissue regeneration and will therefore provide open space for nutrient supply and vascularization for bone tissue formation [41]. Successful cell migration between the scaffolds was also confirmed, in case of assembly (Figure 3g). This ensured continuous cell migration in all three spatial directions and proliferation are possible in any shape assembled ATS configuration and demonstrated that the assembled ATS system can work as a single implant that cells are homogenously spread.

Then, we evaluated *in vitro* mineralization and osteogenesis of ATS scaffold. First, the calcium deposition by ARS staining confirmed the highest mineralization in ST-UP group on both week 1 and week 2 which suggests that the scaffold with higher porosity was able to induce improved cell proliferation, migration of host cells and osteogenic differentiation by providing a macroporous and interconnected structure for cell ingrowth and nutrient supply (Figures 4a and 4b) [4, 41–43]. Similar results were obtained from RT-qPCR, showing that the porous ATS groups exhibited a higher osteogenic gene expression, especially ST-UP (Figures 4c and 4d). As previously mentioned, there are many possible factors that could lead to these results, but two main reasons for strong osteogenic effect of ST-UP is the nanorough surface and interconnected macroporous structure. Especially, for the porous structure, the relationship between pore sizes of the scaffold and osteogenic differentiation of cells has been extensively explored, however, it is still controversial on which specific pore size or geometry is ideal for bone tissue engineering. However, along with other previous studies, the result in this study also suggests that higher porosity and pore sizes of scaffold would result in enhanced osteogenesis and calcium deposition [20, 44, 45]. Furthermore, interconnected porous structure of ATS would result in more stable fixation and better osseointegration with the host bone due to interlocking between the scaffold and the surrounding tissue when it is implanted in physiological system [20, 46].

Finally, we assembled ATS scaffolds to demonstrate the possible implants that can be applied in several medical applications such as spinal fusion surgery (Figure 4e). There has been some studies that showed possible concept of assemblable scaffold. For instance, Subbiah et al., developed 3D-printed hollow synthetic polymer based scaffolds that can be stacked together [47]. However, compared to titanium based scaffold, it is not mechanically stable enough to endure the physiological loading in case of implantation. On the other hand, Lee et al., suggested another strategy by fabricating cylindrical pin and hole structured titanium scaffolds that can be stacked in a two-body combination [48]. However, the scaffolds could be combined in limited way, and the final scaffold combined by inner pin and outer hole could not have interconnected porous structure which made it difficult to have efficient bone ingrowth and nutrient flow. To the best of our knowledge, this is the first study that showed fully assemblable platform that can actually endure the physiological loading as an implant and be assembled to any shape or size that we desired. This would enhance tissue regeneration rate by fitting into the defect and reducing the gap between implant and tissue. In addition, it is also possible for surgeons to control the tissue formation rate of specific region in the defect by choosing specific lattice microarchitectures (NP, SP, UP) or by seeding different cell types/numbers or adding growth factors such as BMP or VEGF in certain ATS units. The biggest advantage of ATS system would be unlimited combinations of ATS assembly, and numerous development based on ATS motif can be achieved by 3D printing with different biomaterials and structures or incorporating drugs or proteins for more precise patient specific tissue regeneration.

## 4. Conclusions

In summary, the concept of the ATS fulfills the needs of patient-specific regenerative medicine by allowing custom, manual assembly and providing stable mechanical properties and positive biological conditions for bone growth in a safe implant. We propose that this ATS has a promising potential to be developed further and extended to further application for bone tissue engineering that requires patient-specific treatment.

## 5. Materials and Methods

### 5.1 Assemblable Titanium Scaffold (ATS) 3D Printing and Assembly

For the 3D printing of ATS, the 3D models were designed using computer-aided design software (Fusion 360, Autodesk), as shown in figure S1. The support structures were created with Magics (V21, Materialise) using contour support. The manufacturing of the parts was done using a selective laser melting (SLM) SLM 250^HL^ system by SLM Solutions GmbH (Lübeck, Germany) with a building platform of 250 × 250 mm^2^, with integrated powder reconditioning and sieving unit. The SLM system used a continuous wave 200 W Ytterbium fiber laser with a wavelength between 1068 and 1095 nm. A layer height of 30 microns was used to print a pure titanium (Ti grade II according to ASTM F67; SLM-Solutions GmbH) with a d_50_ of 41 ± 2 microns (Particle size analysis with Helos/KF + RODOS + VIBRI particle size distribution analysis set up by SympaTec GmbH, Clausthal-Zellerfeld, Germany). The printing parameters were 100 W nominal laser power for the outer contour at a scanning speed of 550 mm/s and 175 W for the inner contour with a scanning speed of 833 mm/s. After the SLM process, the parts were carefully detached manually from the building platform. The support structures were then broken off at perforated support breaking points.

Two different types of scaffold surfaces were used in this study: (1) native SLM, (2) surface textured surfaces (ST) by double acid-etching. For the latter type, 3D-printed SLM scaffolds were treated in a hot mixture of HCl (32%; Fluka): H_2_SO_4_ (95%; J.T. Baker): ultrapure H_2_O (resistivity 18.2 MΩcm, ELGA Purelab Option-Q DV 25) at elevated temperature for 15 min and then rinsed twice with ultrapure water (resistivity 18.2 MΩcm) in an ultrasonic bath. Then, second acid etching was carried out in the same manner as first for the double acid etching treatment.

### 5.2 Finite Element Analysis (FEA)

The mechanical strength of the ATS was demonstrated by FEA under two different loading conditions. A distributed pressure load of 1.25 MPa, the maximum intradiscal pressure reported in other studies [22], was applied on the top surface of the scaffold with the bottom surface fully constrained. Furthermore, a pressure load of 1.25 MPa was applied on the side surface with the opposite side fully constrained. For FEA, titanium alloy, the material of ATS, was assumed to be isotropic and linear elastic with a Poisson ratio of 0.34, a Young’s modulus of 120 GPa and a yield strength of 805 MPa. A commercial FEM solver (NX 12.0) was used to perform the linear static analysis using a mesh with 10-node tetrahedral elements. The mesh resolution was determined after conducting a mesh convergence study. The mechanical behavior was inspected by plotting the Von Misses stress, which should stay well below the yield strength of the material.

To investigate the mechanical strength of the ATS system in case of assemblies, FEA was performed with two or more scaffolds in an assembled configuration. The interface between the scaffolds was assumed to be rigid. This assumption is based on the observation that the scaffolds were tightly fitting after assembly in mechanical testing, therefore displacements between the assembled components can be neglected.

### 5.3 Mechanical test

The ATS groups were tested using a materials testing machine (Schenk RMC 100, Germany) at a displacement speed of 1 mm/min under vertical and lateral compression, respectively. An unconfined quasi-static compression was performed between two parallel smooth plates, and the signals of force and displacement were recorded throughout the experiment.

In addition, to investigate the durability of the ATS in an assembled condition, further dynamic cyclic loading tests were performed using two assembled configurations of the scaffolds. The experiments were carried out on a dynamic material testing machine (Instron E10000, Instron, High Wycombe, UK). Cyclic loading was applied to a maximum force of 150 N at a frequency of 0.05 Hz. 3 cycles of preloading were first performed to ensure that the two scaffolds were tightly coupled together, followed by 5 cycles of compression for property determination.

### 5.4 Scanning electron microscopy (SEM)

Native SLM and ST treated NP, SP and UP ATS were printed, prepared and fixed on metal stubs with carbon tape and coated with platinum sputtering (CCU-010, Safematic). Then, field emission scanning electron microscopy (FE-SEM) (SEM SU5000, Hitachi) was used to capture the macro/microstructure and surface of the scaffolds at 3 kV.

### 5.5 Measurement of surface roughness

First, the surfaces of 3D-printed ATS samples were visualized and captured by confocal laser scanning microscopy (CLSM) (LSM 780 upright, Zeiss, Germany). A 10x magnification lens and 283 x 283 μm^2^ image size were used to capture the Z-stack images of the disc surface. Then, ConfoMap software (Zeiss, Germany) was used for visualization of the surfaces in 3D and measurement of the area roughness parameter, Sq, of the samples. Sq values were calculated according to the ISO 25178.

### 5.6 Microcomputed Topography (Micro-CT) measurement

MicroCT was performed on 3D-printed ST ATS scaffolds using a Scanco μCT 100 instrument (Scanco Medical, Switzerland). Scans were performed at an intensity of 18 W, an energy level of 90 kVp (peak kilovoltage), and an X-ray current of 200 μA. The integration time was set to 140 ms without frame averaging, and with a nominal resolution of 4.9 μm.

### 5.7 Protein adsorption analysis

To measure the protein adsorption capacity of different ATS groups, each scaffold was submerged in 0.1 mg/ml and 1 mg/ml concentrations of BSA solutions for 1 hour at room temperature. After collecting the scaffolds, a Bradford Protein Assay Kit (23200, ThermoFisher) was used to measure the protein adsorbed on the surface of the scaffolds by following the manufacturer’s protocol. For the protein adsorption test with complete medium, the scaffolds were submerged in DMEM/F-12 (31330038, ThermoFisher) with 10% fetal bovine serum (FBS) (26140079, ThermoFisher) and incubated for 1 day. Then, the ATS samples were collected and the protein adsorption assay was carried out in the same manner with a Bradford Protein Assay Kit.

### 5.8 Live & Dead assay and proliferation rate

Mouse pre-osteoblast cells (MC3T3-E1) were obtained from University of Zurich, Zurich, Switzerland. First, 1 × 10^5^ cells were seeded onto the scaffolds. After 1 hour of attachment, cells were cultured in growth medium (GM) composed of MEM α without ascorbic acid (A1049001, Gibco), 10% fetal bovine serum (26140079, Gibco) and 1% antibiotic-antimycotic (15240062, Gibco). After 3 days of culture, cells were stained for 10 min in 0.5 µL/mL calcein-AM and 2 µL/mL ethidium homodimer-1 from the Live/Dead assay kit (L3224, Invitrogen). Then, the cells cultured on scaffolds were fixed in 4% paraformaldehyde (PFA) in PBS for 15 min. The cells on the surface of ATS scaffolds were visualized with confocal laser scanning microscopy (LSM 780 upright, Zeiss) and viability was calculated as the number of live cells divided by the total number of cells. For cell proliferation, PresoBlue assay kit (P50200, ThermoFisher) was utilized by following the standard protocol. First, 1×10^4^ cells were seeded onto the scaffolds, and after 3 hours of incubation for the attachment of cells, the GM was changed to the assay medium containing 10% of PresoBlue solution. After 30 minutes of incubation, the medium changed to the fresh GM, and the assay medium was collected and analyzed by fluorescence spectroscopy (Infinite 200 Pro, Tecan Life Sciences) at excitation wavelength of 560 nm and emission of 590 nm. The same procedure was performed on days 1, 2, 4, 6, 8, 11, 13 and 15, and the percentage of reduction was calculated. The testing groups used for proliferation were ST-NP, ST-SP and ST-UP ATS scaffolds. The number of replicates used in this experiment was four.

### 5.9 Cell attachment and migration

Alexa Fluor 568 Phalloidin (A12380, Thermofisher) and Dapi (62247, Thermofisher) was used to stain actin and cell nuclei by following the standard protocol. After washing with PBS, MC3T3-E1 cultured on scaffolds were fixed in 4% PFA in PBS for 15 min. After being rinsed three times with PBS, cells were permeabilized using 0.1% Triton X-100 in PBS for 15 min and blocked with a 0.1% Triton X-100 in PBS solution with 1% BSA (A2153, Sigma-Aldrich) for 45 minutes. For immunofluorescence staining of actin, cells on scaffolds were stained with fluorescent phalloidin staining solution for 60 minutes. Then, the scaffolds were rinsed at least three times with PBS. After rinsing, for Dapi staining, the samples were stained with the Dapi working solution for 5 minutes and rinsed with PBS to remove excess staining solution. Then, the scaffolds were visualized with confocal laser scanning microscopy (LSM 780 upright, Zeiss). In case of observation for cell migration in assembled ATS scaffolds, first, the cells were seeded onto single ST-UP ATS scaffold. After 3 hours of attachment, cells were cultured with GM for 3 days. Then, the cell laden ATS was assembled with another clean ATS which does not have cells attached. After assembly, the scaffolds were cultured with GM for 1 week, then the dapi staining was carried out in the same manner to observe the cell migration top to bottom ATS scaffold.

### 5.10 *In vitro* osteogenic differentiation

First, osteogenic medium (OM) was made by adding 100 nM of dexamethasone (D4902, Sigma-Aldrich), 10 mM of glycerol-2-phosphate disodium salt hydrate (G9422, Sigma-Aldrich), and 50 µg/mL of L-ascorbic acid (A92902, Sigma-Aldrich) in GM. MC3T3-E1 were seeded on the scaffolds, and cells were cultured with OM. OM was changed every two days, and Alizarin Red S (ARS) staining and RT-qPCR were performed on 7 and 14-day samples. The testing groups used for osteogenic differentiation experiment were ST-NP as a negative control, ST-SP and ST-UP ATS scaffolds.

### 5.11 ARS staining

After 7 and 14 days of osteogenic induction, the ARS kit (0223, Sciencell research) was utilized by following the standard protocol [49]. Cells were fixed in 4 % PFA (281692, Santa Cruz Biotech) for 10 min and washed three times with distilled water. Then, the samples were stained with 2 % ARS solution for 30 min and washed with distilled water until excess staining agents are removed. The amount of mineral content was measured by eluting the ARS with 10% cetylpyridinium chloride (C0732, Sigma-aldrich) and the optical density was measured at OD 570 nm using the microplate reader (Infinite 200 Pro, Tecan Life Sciences). The number of replicates used in this experiment was three.

### 5.12 Real-time Quantitative PCR (RT-qPCR) analysis

Total RNA of osteogenically differentiated pre-osteoblasts at days 7 and 14 were extracted by RNeasy Plus Mini Kit (Qiagen Inc., USA). RT-qPCR was performed to confirm osteogenic gene expression levels by using TaqMan gene expression assays with the following probe/primer combinations: *GAPDH, Mm99999915_g1; ALP, Mm00475834_m1; COL1, Mm00801666_g1; RUNX2, Mm00501578_m1; and OCN, Mm03413826_mH;* (ThermoFisher Scientific, USA). The testing groups used for RT-qPCR were ST-NP as a negative control, ST-SP and ST-UP ATS scaffolds. The number of replicates used in this experiment was three.

### 5.13 Statistical analysis

All experiments were performed at least in triplicate and all data were analyzed as mean ± SD. For statistical analysis, one-way ANOVA was performed followed by Tukey’s post hoc test and statistical significance was considered by p-value: *p<0.05, **p<0.01, and ***p<0.005.

## ASSOCIATED CONTENT

### Supporting Information

**Figure S1.** Design drawings of single NP, SP and UP ATS scaffolds with dimensions.

**Figure S2.** Three-dimensional views and cross-sections of ATS scaffolds.

**Figure S3.** Representative FE-SEM images of SLM and ST treated NP, SP and UP ATS.

**Figure S4.** Average stiffness of single ATS under vertical and lateral compression. Open bar indicate SLM, hatched bars represent ST samples.

**Figure S5.** Representative force-displacement curve for assembled ATS scaffolds (assembled longitudinally with two units) under cyclic compressive loading.

**Figure S6.** Representative SEM images of cell attached on ATS scaffolds.

## AUTHOR INFORMATION

**Corresponding Author**

*

## Acknowledgments

This project has received funding from the European Union’s Horizon 2020 research and innovation programme under the Marie Skłodowska-Curie grant agreement No 812765. The authors also acknowledge use of the Scientific Center for Optical and Electron Microscopy (ScopeM) of ETH Zurich.

## Data availability

The main data supporting the results in this study are available within the paper and its Supporting Information. All raw and analyzed datasets generated during the study are available from the corresponding author on request.

