## Supporting Information for "3D-Printed LEGO^®^-inspired Titanium Scaffolds for Patient-Specific Regenerative Medicine"

Hönggerbergring 64, HPP O24, 8093 Zürich, Switzerland

### Supporting Information Captions

**Figure S1.** Design drawings of single NP, SP and UP ATS scaffolds with dimensions.

**Figure S2.** Three-dimensional views and cross-sections of ATS scaffolds.

**Figure S3.** Representative FE-SEM images of SLM and ST treated NP, SP and UP ATS.

**Figure S4.** Average stiffness of single ATS under vertical and lateral compression. Open bar indicate SLM, hatched bars represent ST samples.

**Figure S5.** Representative force-displacement curve for assembled ATS scaffolds (assembled longitudinally with two units) under cyclic compressive loading.

**Figure S6.** Representative SEM images of cell attached on ATS scaffolds.

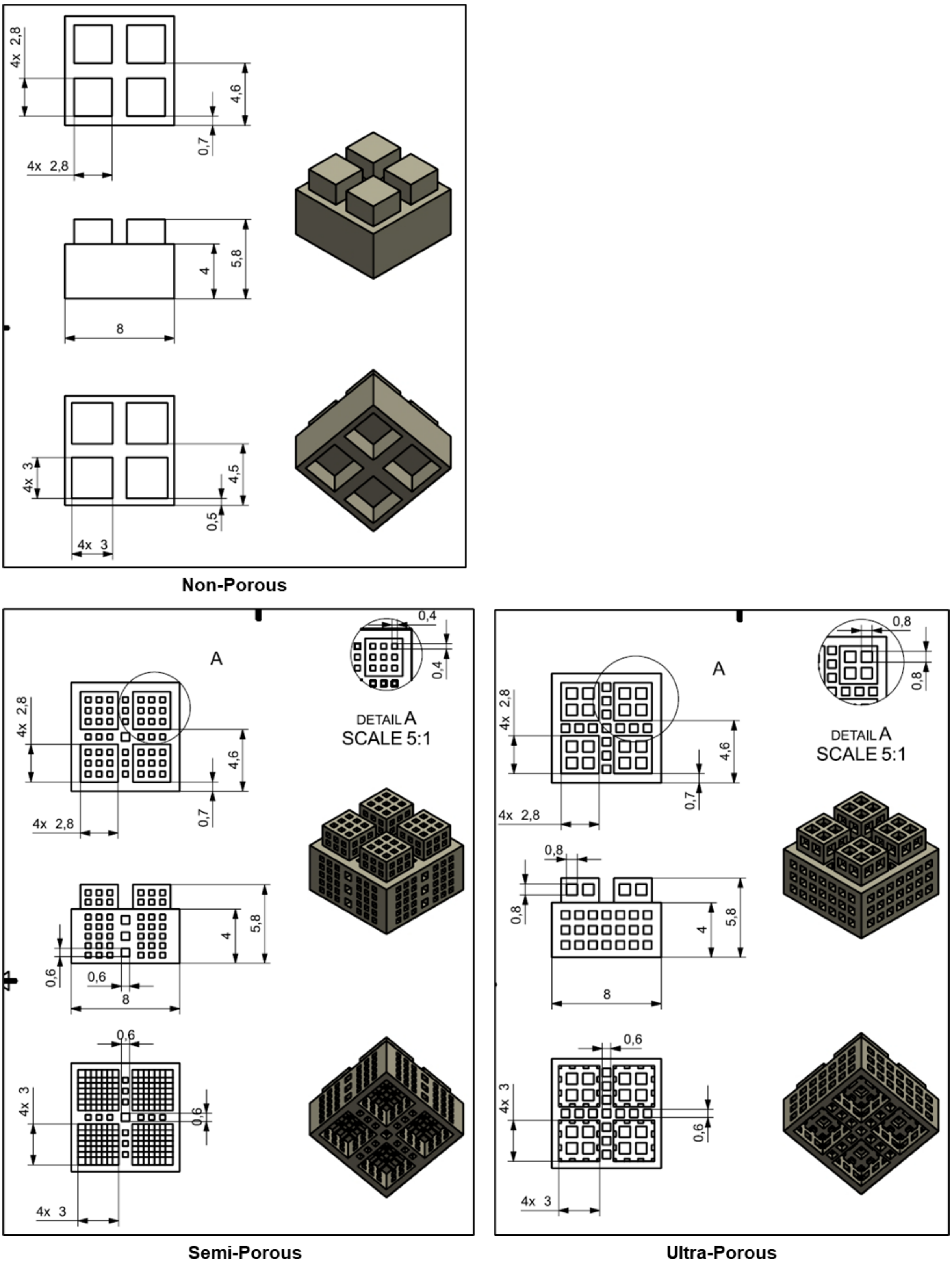


**Figure S1.** Design drawings of single NP, SP and UP ATS scaffolds with dimensions.

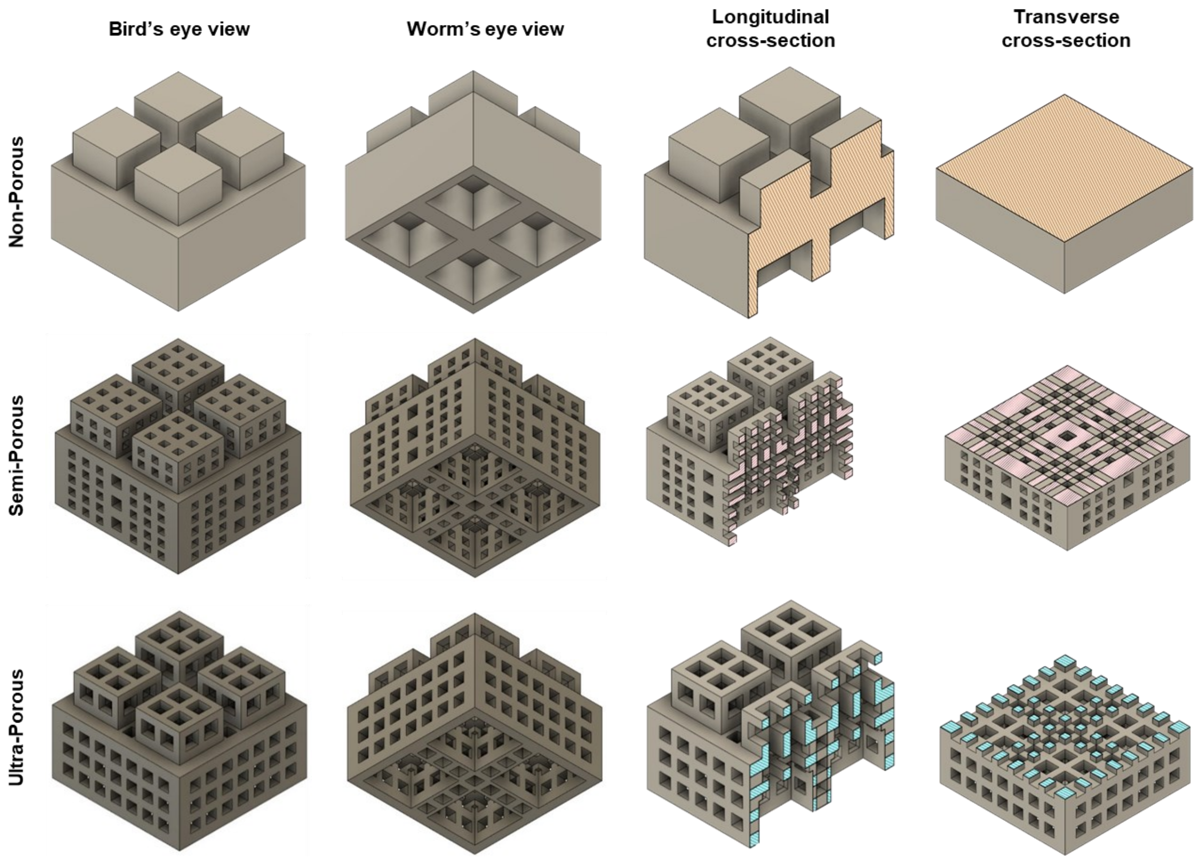


**Figure S2.** Three-dimensional views and cross-sections of ATS scaffolds.

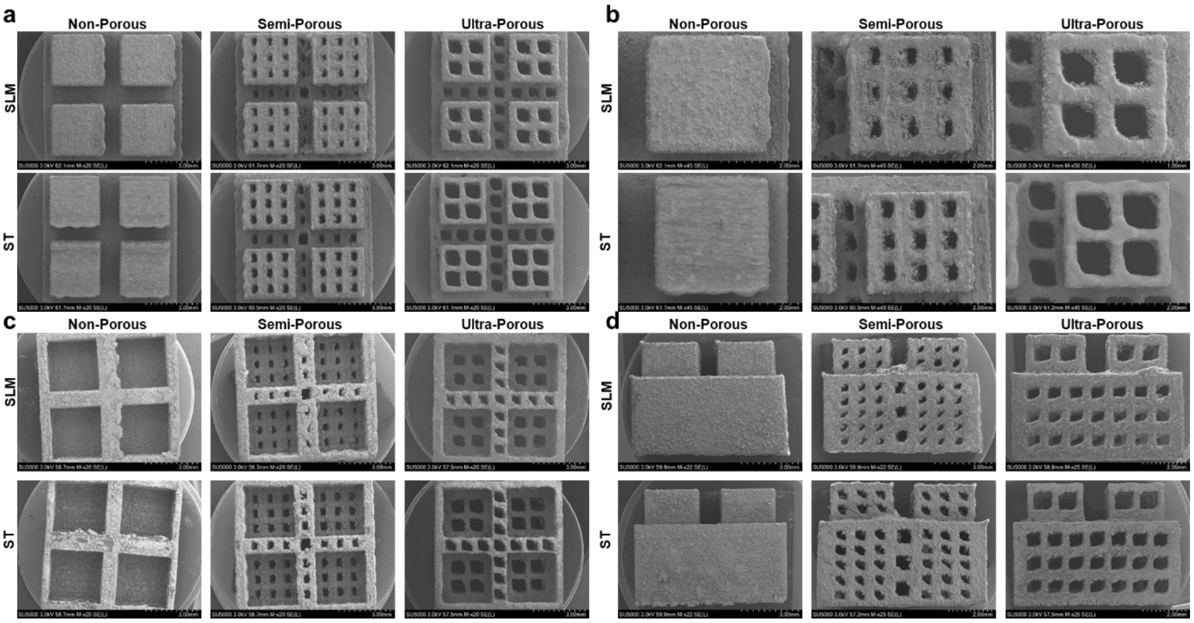


**Figure S3.** Representative FE-SEM images of SLM and ST treated NP, SP and UP ATS. Each group of FE-SEM images was taken from different perspectives which are (a) top view, (b) magnified image of (a), (c) bottom view and (d) side view.

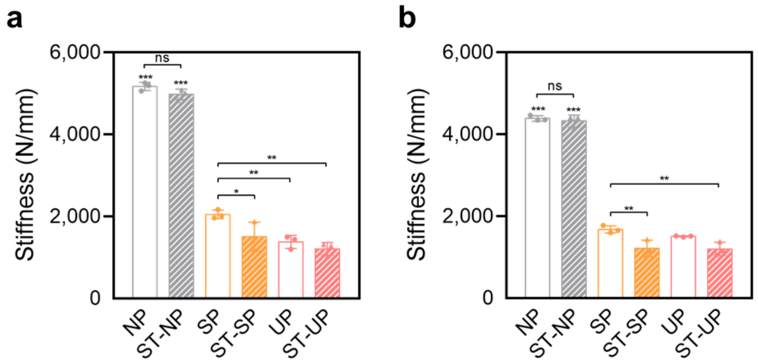


**Figure S4.** Average stiffness of single ATS under (a) vertical and (b) lateral compression. Open bar indicate SLM, hatched bars represent ST samples.

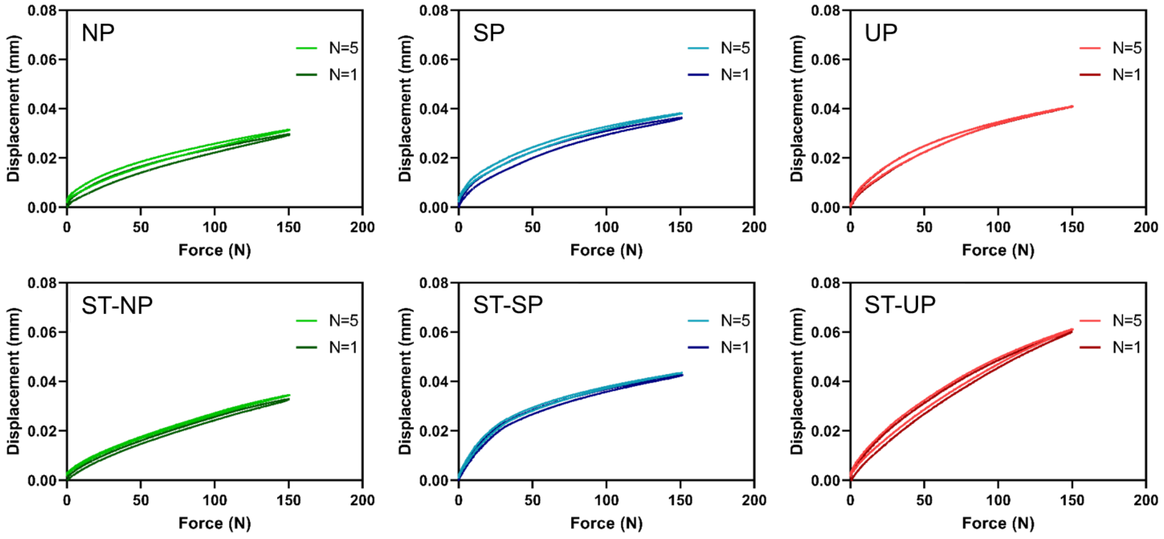


**Figure S5.** Representative force-displacement curve for assembled ATS scaffolds (assembled longitudinally with two units) under cyclic compressive loading.

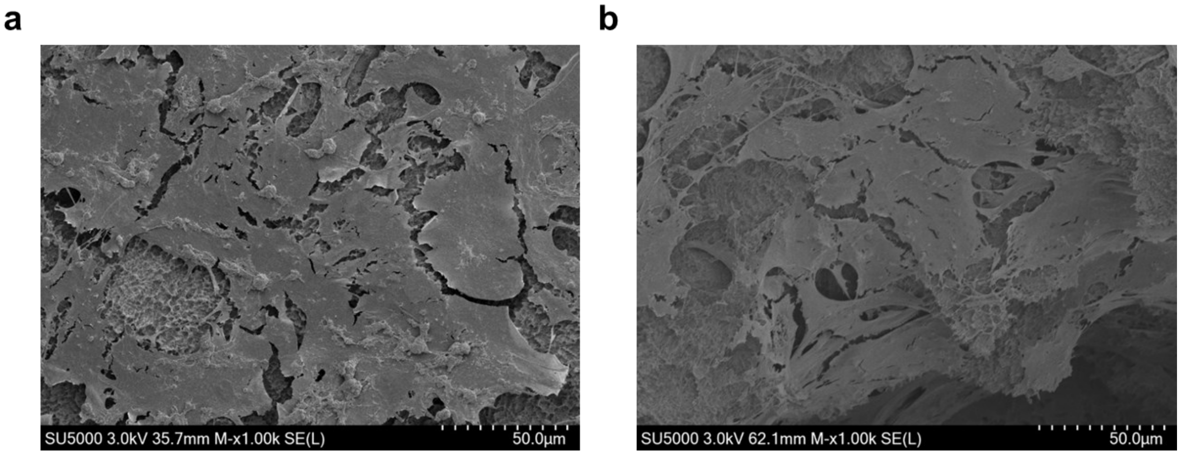


**Figure S6.** Representative FE-SEM images of cell attached on ATS scaffolds. FE-SEM images of cell attachment on (a) ST-NP and (b) ST-UP scaffolds were taken after seeding and culturing pre-osteoblasts for 3 days on the scaffolds.
